# A Modified Comprehensive Grading System for Murine Knee Osteoarthritis: Scoring the Whole Joint as an Organ

**DOI:** 10.1101/2021.05.05.442864

**Authors:** Caleb W. Grote, Matthew J. Mackay, Xiangliang Liu, Qinghua Lu, Jinxi Wang

**Author notes:** Correspondence to: Jinxi Wang, MD, PhD, Department of Orthopedic Surgery University of Kansas Medical Center, 3901 Rainbow Boulevard, MS #3017, Kansas City, KS 66160, USA.

## Abstract

**Objective:** This study aimed to develop a comprehensive but easy to apply histologic grading system to score osteoarthritic changes in the whole knee joint for both spontaneous and posttraumatic osteoarthritis (OA) mouse models.

**Methods:** The new OA grading scheme was developed based on extensive literature review and the authors’ experience in mouse OA models with relatively long periods of observation (up to 24 months of age or 24-week post-surgery). Semi-quantitative assessments of the histopathologic OA changes were applied to all four quadrants of the knee. Grading elements per quadrant were defined as follows: cartilage (0-7) including three new grading elements for early- and late-stage OA, respectively; osteophyte (0-2) covering chondro-osteophytes in both outer and inner joint margins; subchondral bone (0-2) containing subchondral bone thickening and destruction; synovitis (0-2) comprised of both synovial plica and intercondylar notch; and peri-articular tissues (0-2) highlighting ectopic chondrogenesis and ossification in the knee capsule, ligament, and musculature.

**Results:** Statistical analyses showed that the new grading system had high intra- and inter-observer reproducibility (Pearson’s correlation coefficients r >0.9) for both experienced and novice scorers. Sensitivity and reliability analyses confirmed the ability of the new system to detect minimal OA progression between two timepoints with a two-week interval and to accurately identify tissue-specific OA severity within the knee joint.

**Conclusions:** The comprehensive histologic grading system presented here covers all-stage osteoarthritic changes in all major knee joint tissues of mice, which enable us to score OA severity for the whole joint reproducibly and accurately without software-assisted time-consuming measurements.

## Introduction

Osteoarthritis (OA) is one of the leading causes of chronic disability in the United States and has a significant socioeconomic burden accounting for $185 billion dollars each year in medical cost and lost work ^1; 2^. Accurate and reproducible histologic assessment of OA severity in animal models is critical in studies evaluating the efficacy of therapeutic agents designed to prevent or attenuate disease severity.

While multiple animal models of OA have been developed for scientific studies, mouse models remain the most popular due to relatively low cost, easy breeding, modifiable genetics, reliable interventions and predictable outcomes ^3–5^. Previously reported animal models of OA have substantially improved our understanding of OA pathogenesis. However, most OA grading systems are not standardized, making comparative studies and communication among researchers difficult. Furthermore, although the subchondral bone, cartilage, synovium, capsule, and ligaments may all play a role in OA development and progression ^6; 7^, the most widely used existing OA grading systems focus almost exclusively on cartilage changes ^8–10^. To date, few OA grading systems have characterized the pathological changes of multiple joint tissues and they often require complex time-consuming measurements. An easily executed, comprehensive OA grading system is lacking. Developing a universal mouse OA grading system that incorporates all known aspects of OA pathology has several potential benefits: 1) Improving communication among researchers by standardizing results, 2) identifying early, potentially modifiable changes in joint tissues prior to structural damage or articular destruction, 3) further elucidating OA pathogenesis, and 4) scoring OA severity for the entire joint as an organ.

The objective of this study was to develop a more comprehensive but less time-consuming histologic OA grading system that can be applied universally to murine models of post-traumatic OA (PTOA) and spontaneous OA. The novel modified OA grading system reported here was developed based on our extensive literature review and decades of OA histology experience with a spontaneous OA model established in our own laboratory ^11–14^ and a PTOA model induced by surgical destabilization of the medial meniscus (DMM) as described by Glasson et al.^15^. The modified grading system includes analysis of early cartilage degradation and more severe cartilage lesions that were not included in the previously reported grading systems, osteophyte formation, subchondral bone changes, synovitis, and ectopic chondrogenesis with endochondral ossification in the peri-articular soft tissues. This reproducible semi-quantitative grading system does not require complicated calculations or laborious and time-consuming measurements. In addition, it fulfills the principles of an ideal OA grading system as defined by the OARSI study group including simplicity, utility, scalability, extendibility, and comparability ^9^.

## Materials and Methods

### Murine models of osteoarthritis

The development of this histological grading scheme was mainly based on our OA histology experience with both spontaneous and Posttraumatic OA. Time-dependent spontaneous OA changes in the Nfat1-deficient (*Nfat1*^−/−^ and *Nfat1*^+/−^) and aged wild-type (WT) mice were observed up to 24 months of age^11; 13^. Posttraumatic OA in mice induced by destabilization of the medial meniscus (DMM) or anterior cruciate ligament transection (ACLT) with or without treatments were examined at 2, 4, 8, 16, and 24 weeks post-surgery depending on the specific research purposes (unpublished results). At least five animals per stain/gender were evaluated at each time point. All animal procedures were performed with the approval of the Institutional Animal Care and Use Committee in compliance with all federal and state laws and regulations.

### Tissue section preparation and histopathologic analysis

Mouse knee joint tissue samples were fixed in 2% paraformaldehyde, decalcified in 25% formic acid, embedded in paraffin, and sectioned coronally to examine both the medial and lateral compartments. Tissue sections at 5-μm-thick were obtained every 80-100μm across the entire joint. Safranin-O stain was utilized to specifically identify cartilage cells and matrices. Hematoxylin-Eosin (H&E) stains were used to examine the structure of bone and periarticular tissues. Histologic images were acquired with a ZEISS-Axioskop microscope equipped with a digital camera at 25X, 50X, 100X, and 200X. Generally, 25X and 50X images were sufficient for grading osteoarthritic cartilage lesions, chondro-osteophyte formation, subchondral bone changes, and remarkable periarticular soft tissue alterations; 100X and 200X images can be used for cellular analysis of articular cartilage and synovium. Histopathologic analysis was conducted as described previously^11–14^.

### Literature review to identify the necessity of modifications

A comprehensive review of published articles (PubMed, 1960-2020) on OA histology was conducted, mainly focused on the application to rodent OA models. The literature review found that although most histopathologic features of OA had been described in the existing grading systems, many important OA characteristics seen in murine models of OA were still not included. More importantly, most widely used existing OA grading systems focus almost exclusively on cartilage changes ^9; 10^. Although there are a few articles describing OA grading of multiple joint tissues, most assessment methods are complex and time-consuming ^16; 17^. For these reasons, we proposed a novel comprehensive knee OA grading scheme that covers more joint tissues with relatively simple and less time-consuming scoring methods. A comparison of the existing grading systems and our newly developed grading scheme are summarized in **Table 1**.

**Table 1.**
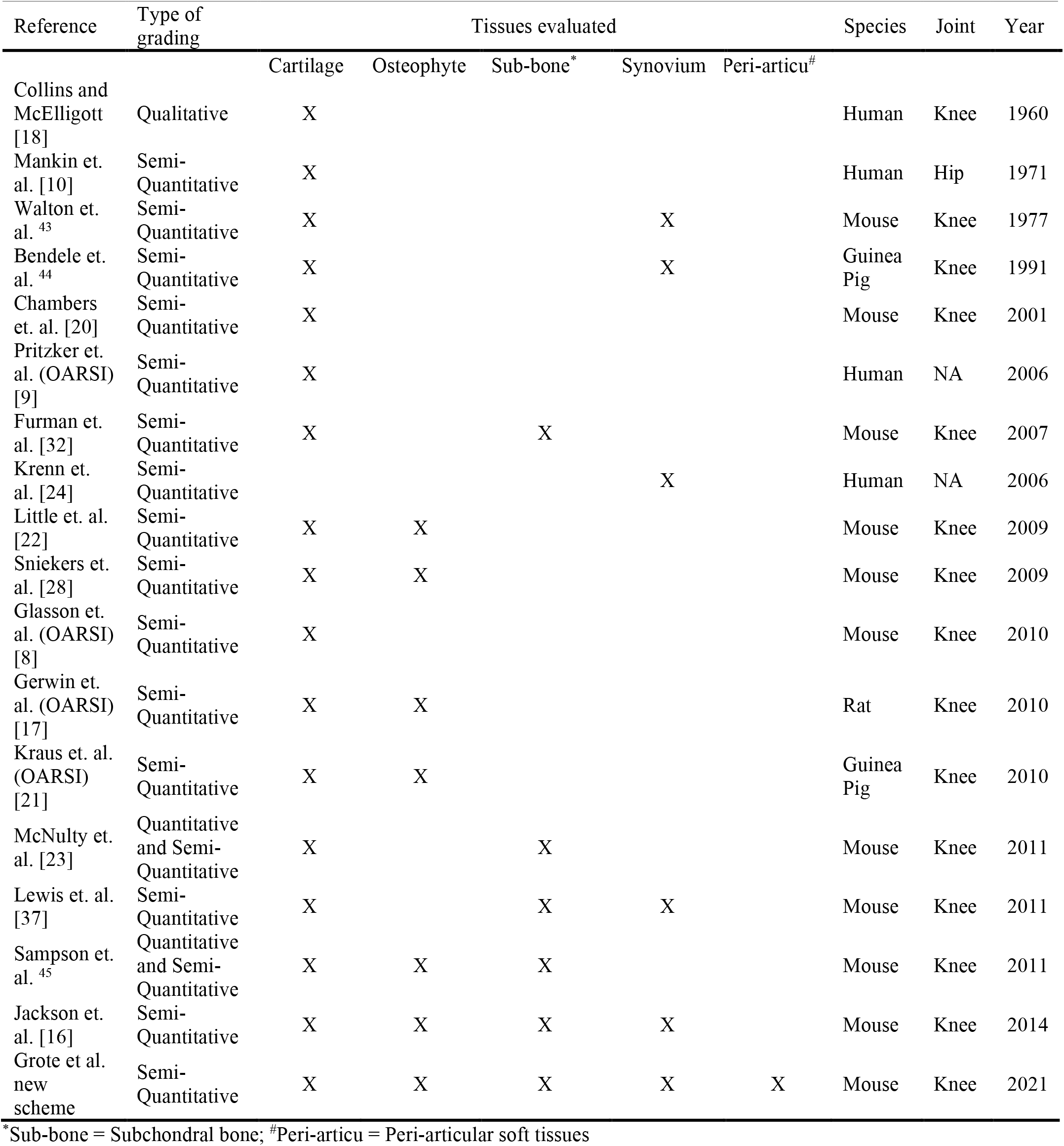
Comparison of commonly referenced OA grading systems with proposed new scoring scheme

The widely used existing OA grading systems and justifications for developing our new knee OA grading system are described below.

#### Articular cartilage lesion

Following the pioneering grading system reported in 1960 by Collins and McElligott ^18^, the Mankin Grading System or Histologic Histochemical Grading System (HHGS) was published in 1971 for the evaluation of human hip OA ^10^ and has later been adapted to histologic evaluation of multiple animal models of OA, including mice. Mankin scoring focuses on cartilage changes including cartilage structure (0-6), chondrocyte cellularity (0-3), safranin-O staining (0-4), and tidemark integrity (0-1). However, concerns about the accuracy and reliability of the Mankin System have been raised, particularly for the thin cartilage of mice ^19^. Secondary to these concerns and in an effort to standardize OA grading, the Osteoarthritis Research Society International (OARSI) formed a working group that published a new OA grading system in 2006 ^9^. The OARSI grading system was focused on cartilage including grades 0-6 based on cartilage surface integrity, surface discontinuity, cartilage clefts/fissures, erosions, denudation, and deformation. This system was developed mainly based on human OA pathology but has also been used for rodent OA.

The OARSI histopathology initiative focusing on mouse models of OA was published in 2010 ^8^. This mouse OA grading system was modified from a previous grading system by Chambers et. al. ^20^ and mainly focused on cartilage based on the depth and percent of surface area of cartilage lesions (score 0-6), though 0-3 scoring parameters for each of osteophyte, subchondral bone and synovitis were proposed without clear definitions. In addition to those widely used grading systems for OA cartilage, other OARSI histopathology initiatives and cartilage/synovium OA grading systems have also been reported and utilized ^16; 17; 21–24^.

Our new cartilage scoring method was modified from the OARSI ^8^ and Chambers et. al. ^20; 25^ grading systems. We propose grades 0-7 per quadrant including grade 1.5 to highlight the loss of surface lamina for the early phase of OA and simplify the scoring process by compressing their grades 3-6, changing the criteria to <34%, 34-67%, and > 67% cartilage loss as opposed to increments of 25%. In addition, two new grades for lesions beyond the calcified cartilage layer to the level of subchondral bone were added to accommodate more severe models, which were frequently seen in mouse models of spontaneous OA (late stages) and posttraumatic OA (16 weeks or longer post-surgery). The representative histologic images demonstrating our novel cartilage scoring elements are presented in **Figure 1**.

**Figure 1.**
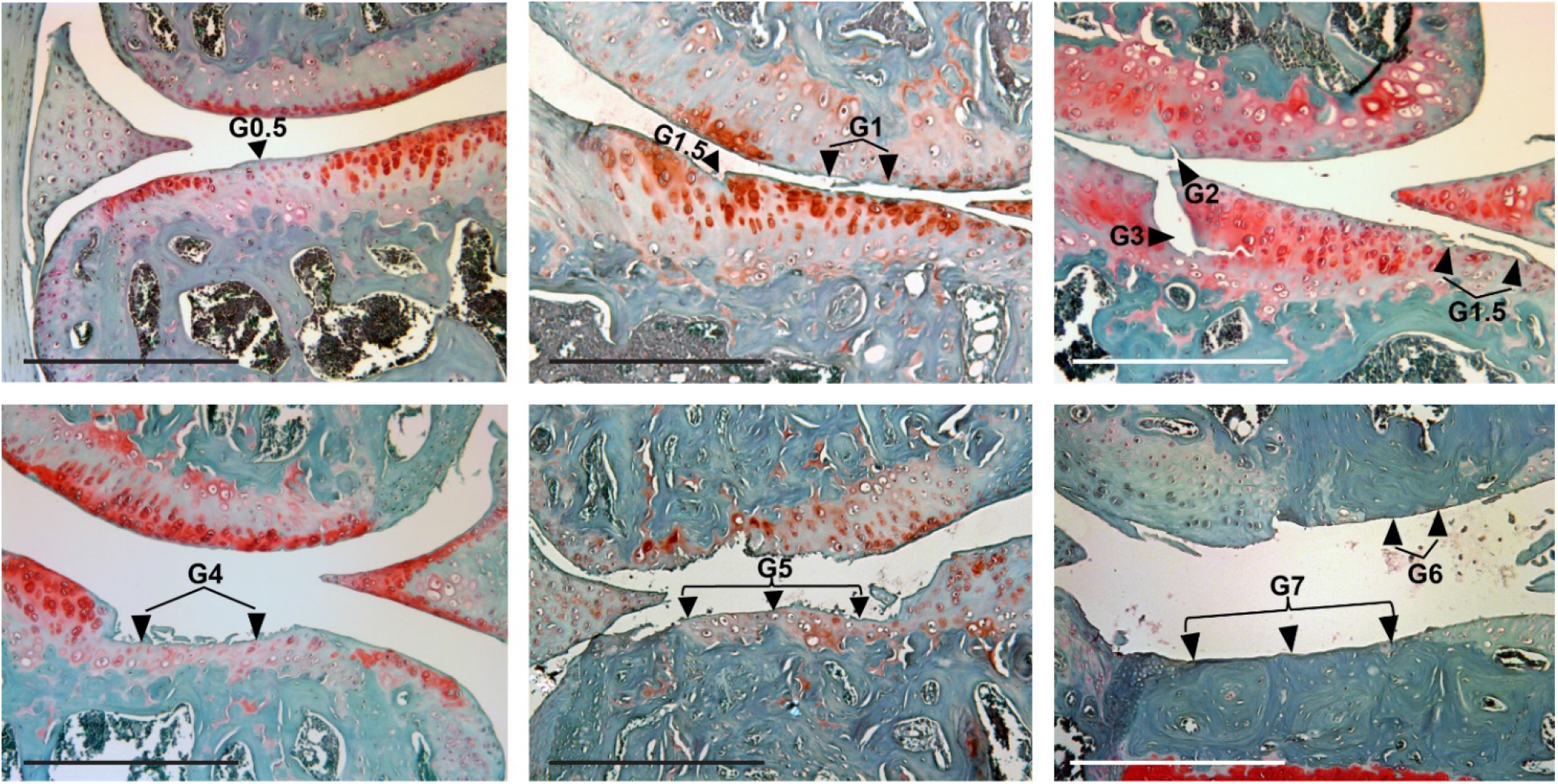
Photomicrographs showing osteoarthritic cartilage lesions in the femoral condyle (upper) and tibia plateau (lower) of each micrograph. Arrowheads point to specific cartilage lesions with various OA cartilage grades as described in Table 1A. G0.5 = grade 0.5, G1 = grade 1, G1.5 = grade 1.5, G2 = grade 2, G3 = grade 3, G4 = grade 4, G5 = grade 5, G6 = grade 6, and G7 = grade 7. Safranin-O and fast green stains, counterstained with hematoxylin. Scale bar = 100 µm.

#### Osteophyte formation

Previous grading of osteophytes in murine models of OA has not been standardized, though osteophyte scoring methods were included for rats ^17^ and guinea pigs ^21^ as part of the OARSI histopathology initiatives. These osteophyte grading methods are mainly based on sizes of osteophytes. Additional grading systems for osteophytes have been reported ^22; 26–29^. Little et. al. utilized a semi-qualitative histology system based on osteophyte size in relation to the depth of the surrounding articular cartilage (0-3) and on osteophyte maturity ranging from primarily cartilaginous to primarily mature bone (0-3) ^22^. A similar grading system based on osteophyte maturity was also described by Kamekura et. al. ^26^. Osteophytes have also been graded with a categorical system of present or not to calculate the overall incidence ^27; 28^. The use of micro-CT for evaluation of osteophytes has also been reported with more detailed quantitative data ^29^.

Osteophytes in the proposed new grading scheme are graded 0-2 for each quadrant of the knee joint based on presence of chondro-osteophytes at the inner and/or outer joint margins (Grade 0 = no osteophytes, Grade 1 = osteophyte at either inner or outer joint margin, Grade 2 = osteophyte at both inner and outer joint margins). This highlights the number and anatomic location of osteophytes and eliminates the need for measuring size and can easily and quickly be scored (**Figure 2A**).

**Figure 2.**
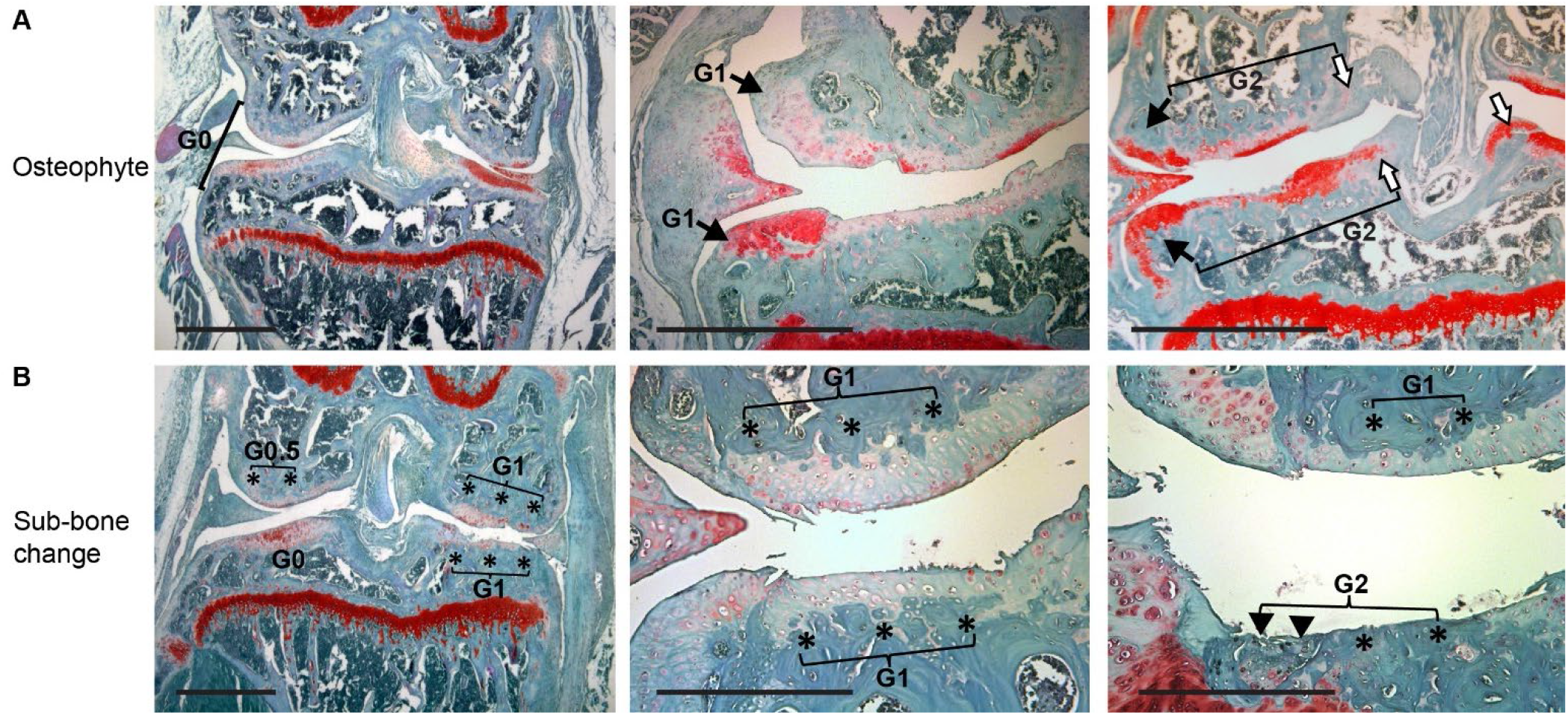
(**A**) Photomicrographs showing chondro-osteophyte formation in the joint margin of the femoral condyle (upper portion of each graph) and tibial plateau (lower). Arrows point to chondro-osteophytes in the outer joint margins; open arrows indicate chondro-osteophytes in the inner joint margins. As described in Table 1B, G0 = no osteophytes, G1 = osteophyte grade 1, G2 = osteophyte grade 2. (**B**) Photomicrographs showing subchondral bone (Sub-bone) changes in the femoral condyle (upper) and tibial plateau (lower). * indicates thickened subchondral bone; arrowheads point to the area of subchondral bone destruction/defect which is filled with fibrocartilage-like tissue. G0 = no Sub-bone changes, G1 = Sub-bone grade 1, G2 = Sub-bone grade 2. Safranin-O and fast green stains, counterstained with hematoxylin. Scale bar = 100 µm.

#### Subchondral bone change

Subchondral bone plays an integral role in cartilage support and is a key component of OA pathophysiology ^30^. Known radiographic changes of subchondral bone in human OA include sclerosis and cyst formation, but only recently has a histological grading system in human knee joints been proposed ^31^. Subchondral bone thickness in OA mice has been described using a modified Mankin score and reported in relation to the overlying articular cartilage (0-2) ^32^, although this can be problematic if concomitant cartilage loss is present. In the rat, damage to the calcified cartilage layer and subchondral bone is graded together as a supplemental OARSI score, scores 0-5 based on multiple factors including: tidemark basophilia, calcified cartilage collapse, marrow mesenchymal changes, subchondral bone thickening, and articular cartilage collapse ^17^. Grading of guinea pig subchondral bone changes has been extensively studied with micro-CT ^21; 33^ and this has been applied to mouse models as well ^34^. Subchondral bone thickness and area has been quantified with histological slides using computer histomorphometry software ^23^. Those grading changes in subchondral bone either rely on additional studies (micro-CT) or time-intensive measurements for quantitative analysis or unreliable cartilage depth as internal controls.

In the proposed new system, subchondral bone was graded as 0, 0.5, 1, and 2 per quadrant, where grade 0 = no changes, grade 0.5 = focalized subchondral bone thickening with bone marrow cavity in some subchondral area, grade 1 = extensive subchondral bone thickening with essential obliteration of marrow cavity, and grade 2 = subchondral bone thickening with cortical bone destruction or cyst formation (**Figure 2B**). Subchondral bone thickening was defined as a visual increase in bone substance of approximately >1.5 times the normal subchondral bone. While seemingly subjective, multiple direct measurements including imaging with a micrometer on slides and software-assisted quantitative area calculation were performed and confirmed that this method proved to be reliable and easily adapted even by novice reviewers.

#### Synovitis

The involvement of synovial inflammation in rheumatoid arthritis is well known, but low-grade synovitis also plays a role in OA ^35; 36^ and is a key component to the “joint as an organ” approach to OA pathogenesis. A histological grading system for human synovitis ^24^ was recently applied to mouse PTOA models, which demonstrated increasing synovitis with increased severity of intra-articular lesions ^37^. This is a similar system as reported by the OARSI for rats and guinea pigs ^17; 21^, but the OARSI recommendations for grading synovitis in mice are not clearly defined ^8^.

Synovitis was graded as 0, 0.5, 1, and 2 in our new grading system for a complete evaluation of various locations of the joint. Grade 0 represents no evidence of synovitis; grade 0.5 demonstrates focalized thickening at either the side synovial plica (fold) adjacent to the joint capsule or the intercondylar notch (for lower femur) and intercondylar eminence (for upper tibia); grade 1 represents extensive thickening of the synovium including both synovial lining (intima) and subintimal stroma tissue that can be observed at either the side synovial plica or intercondylar notch/intercondylar eminence; and grade 2 demonstrates synovial thickening that can be observed in both areas (**Figure 3**). Thickening was defined as visual increase in synovial thickness approximately >1.5 times the normal synovial thickness, which was compared with direct measurements as described in the subchondral bone section and proved to be easily identifiable and reliable. This highlights the presence of low grade/mild synovitis in OA joints in various anatomic locations, while removing the need for time-consuming cellular analysis and cell-layer counting.

**Figure 3.**
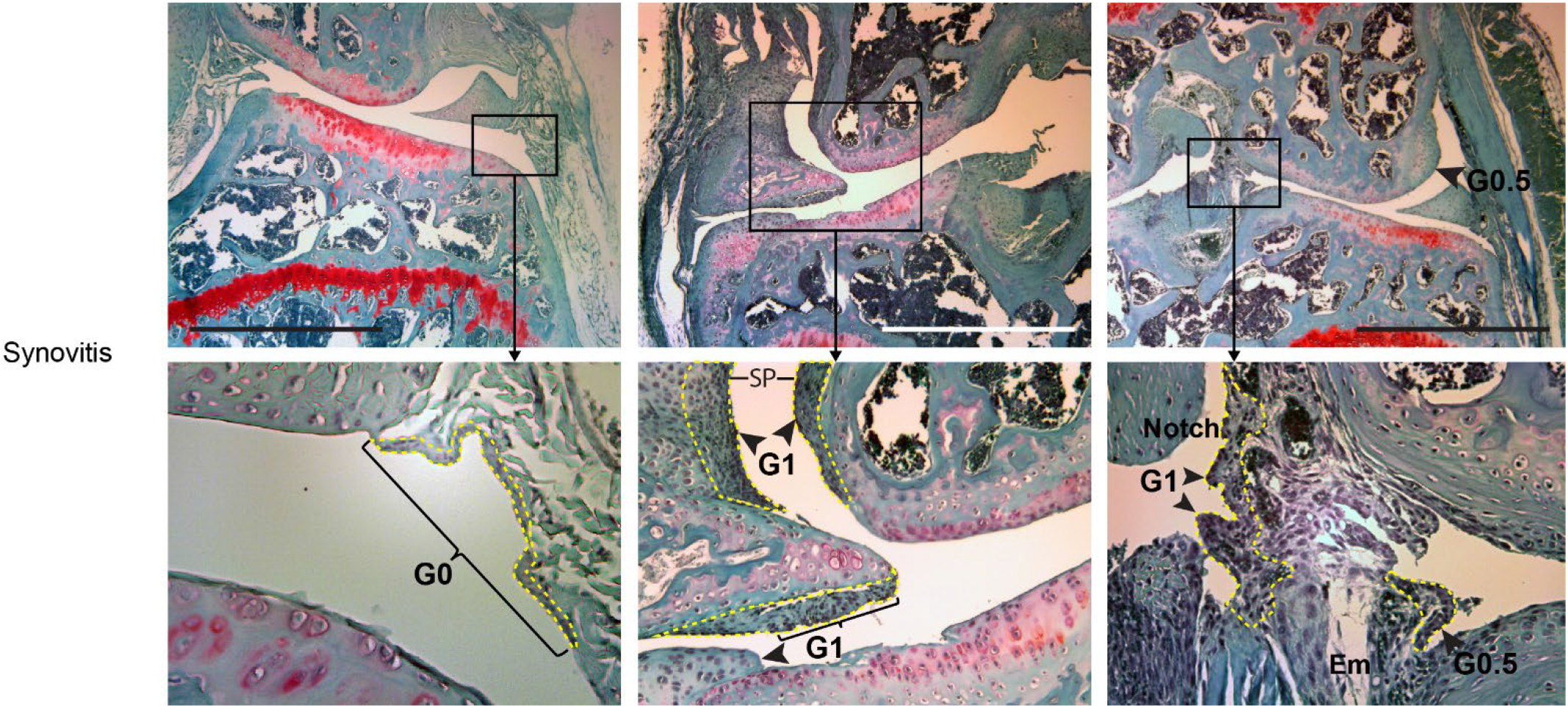
Photomicrographs showing different grades of synovitis at low (upper panels) and higher (lower panels) magnifications. **Left**: Normal synovium (grade 0/G0) with a single layer of synovial lining cells (outlined by a dotted yellow line) and scattering cells in the subintimal connective tissue. **Middle**: Extensive synovitis (grade 1/G1) in the side synovial plica (SP) around the femoral condyle (upper portion) as well as the meniscus and joint margin (lower portion). **Right**: Extensive synovitis (G1) in the femoral intercondylar notch (Notch) with focalized synovitis (G0.5) in the side synovial plica and the tibial intercondylar eminence (Em). For both Middle and Right panels, arrowheads point to thickened synovium with multi-layer synovial lining cells (outlined by dotted yellow lines) with cell infiltration in the subintimal connective tissue. Scale bar = 100 µm.

#### Ectopic chondrogenesis and endochondral ossification in periarticular soft tissues

Calcification of cartilage and periarticular structures has been described as one of the osteoarthritic characteristics in humans ^38; 39^. Ectopic chondrogenesis with endochondral ossification has also been demonstrated in ACLT-induced mouse models of PTOA ^15^. Close evaluation of mouse OA models used in our laboratory further demonstrated periarticular chondrogenesis and ossification in the joint capsule as well as surrounding ligament and musculature, distinct from chondro-osteophyte formation at the joint margins. These histologic changes have not been included in any existing grading systems.

Ectopic chondrogenesis with or without endochondral ossification was added to the new grading system. Grade 0 represents no abnormalities. Grade 1 demonstrates ectopic chondrogenesis and/or ossification in the synovium and/or capsule, and grade 2 has the change extending into the surrounding ligament and/or musculature (**Figure 4**).

**Figure 4:**
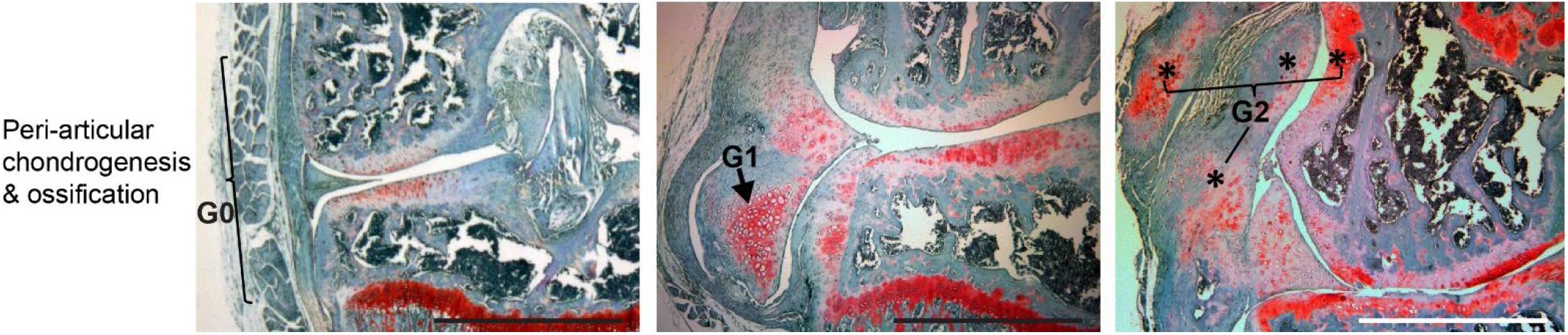
Photomicrographs showing different grades of ectopic peri-articular chondrogenesis and endochondral ossification. **Left panel** (**G0**): No ectopic peri-articular chondrogenesis or ossification. **Middle panel** (**G1**): Ectopic peri-articular chondrogenesis and ossification is limited in the synovium-capsule. **Right panel** (**G2**): Ectopic peri-articular chondrogenesis and ossification is seen in both the synovium-capsule and the peri-articular ligament. Scale bar = 100 µm.

#### Terminology of the new OA scoring system

##### Anatomic term

Knee joint = stifle joint (for veterinary science)

##### Maximum score

Proposed scores/grades for cartilage (0-7), osteophytes (0-2), subchondral bone (0-2), synovitis (0-2), ectopic endochondral ossification (0-2) were set for each quadrant (medial femoral condyle, medial tibial plateau, lateral femoral condyle, and lateral tibial plateau) of the knee joint. This resulted in a proposed maximum score per quadrant at 15 and maximum score per joint at 60 (**Table 2A–B**).

**Table 2A.** Modified comprehensive knee OA grading system: Cartilage lesions.

| Grade* | Articular cartilage lesion |
| --- | --- |
| 0 | None |
| 0.5 | Loss of Safranin-O or other proteoglycan staining without structural changes |
| 1 | Focalized surface abrasion or fibrillation without loss of cartilage |
| 1.5 | Surface fibrillation within the superficial layer with discontinuity of cartilage surface lamina |
| 2 | Vertical clefts/cartilage loss down to the layer immediately below the superficial layer |
| 3 | Vertical clefts/cartilage loss to the calcified cartilage extending to <34% of the articular surface |
| 4 | Vertical clefts/cartilage loss to the calcified cartilage extending to 34-67% of the articular surface |
| 5 | Vertical clefts/cartilage loss to the calcified cartilage extending to >67% of the articular surface |
| 6 | Cartilage loss to the subchondral bone extending to <50% of the articular surface |
| 7 | Cartilage loss to the subchondral bone extending to >50% of the articular surface |

\*Grade/score number is set for each quadrant of the knee joint.

**Table 2B.** Modified comprehensive knee OA grading system: Other changes.

| Grade* | Chondro-osteophyte | Score* | Subchondral bone change | Score* | Synovitis | Score* | Peri-articular chondrogenesis |
| --- | --- | --- | --- | --- | --- | --- | --- |
| 0 | None | 0 | None | 0 | None | 0 | None |
|  |  | 0.5 | Focalized Subchondral bone thickening<sup>#</sup> | 0.5 | Focalized synovial thickening<sup>#</sup> |  |  |
| 1 | Osteophyte present in either outer or inner joint margin | 1 | Extensive subchondral bone thickening | 1 | Extensive thickening in either side plica or intercondylar notch/eminence | 1 | Ectopic chondrogenesis and/or ossification in synovium-capsule |
| 2 | Osteophyte present in both inner and outer joint margins | 2 | Subchondral bone thickening with bone plate destruction, bone cyst, or chondrogenesis | 2 | Extensive thickening in both side plica and inter-condylar notch/eminence | 2 | Ectopic chondrogenesis and/or ossification in both synovium-capsule and ligament/muscle |

\*Grade/score number is set for each quadrant of the knee joint.
<sup>#</sup>Criterion of subchondral bone thickening or synovial thickening is set for >1.5-fold (visual estimate) over the same normal tissue.

##### Quadrantal OA score and whole joint OA score

Quadrantal OA score is referred to as an actual OA score or an averaged OA score for a quadrant of the knee joint from multiple observers. Whole joint OA score (or joint OA score) is referred to as a total/summed score for the entire knee joint, including scores from all four quadrants. Scores also can be reported as or converted to an average of anatomic location subsets (e.g. medial femoral condyle and medial tibial plateau quadrants) or tissue subsets (e.g. cartilage, synovium, and subchondral bone, etc.) from multiple joints/images for statistical analysis.

### Semi-quantitative histologic OA scoring

Representative tissue histologic images with NFAT1 deficiency-mediated spontaneous and surgery-induced posttraumatic knee OA were utilized for semi-quantitative scoring to test the reproducibility and accuracy of the proposed new OA scoring scheme. Midcoronal sections with intact knee tissue structures covering articular cartilage, subchondral bone, joint margin, synovium, joint capsule, and surrounding ligament and musculature were selected for scoring. A set of images from 20 different tissue sections with various levels of knee OA were scored twice with one week apart to obtain intra-observer variability from the same observer. To assess the inter-observer variability as well as the ease of use of the scoring scheme, the images were scored by three independent observers (scorers), including an experienced observer (JW, Observer a), a trained observer (CG, Observer b), and a novice observer (MM, Observer c) who was familiar with mouse knee structures but had no previous experience in OA histology. Another set of images from an additional 20 tissue sections selected from 2 and 4 weeks following DMM surgery (6 slides per group/time point) were scored by all 3 observers who were blinded to the group information to examine the sensitivity and accuracy of the new OA scoring scheme for OA progression between the two time points.

### Statistics

Quadrantal OA scores or whole joint OA scores of representative tissue sections from three observers at each time point were averaged and presented as the mean ± standard deviation or 95% confidence intervals from at least six independent repeats/group. Inter-and intra-observer variability or reproducibility was determined by Pearson’s correlation coefficient analyses. Significant differences between study groups were determined by either Student’s *t*-tests or ANOVA. Statistical analysis was conducted using Microsoft Excel software (Version 2021). A p value of less than 0.05 was considered statistically significant.

## Results

### Validation of inter- and intra-observer reproducibility

Histologic images from twenty tissue slides with mild, moderate, or severe OA pathology, which were derived from both spontaneous and posttraumatic knee OA in mice, were utilized for inter-and intra-observer correlation coefficient analysis to validate the reproducibility of the new grading scheme. The results demonstrated high reproducibility with inter-observer correlation coefficients of >0.90 for both first and second measurement of the whole joint scores across the experienced, trained, and novice observers (**Figure 5A**). Intra-observer variability analysis of the same set of histologic images also showed high reproducibility between the first and second measurement scores for each of the three observers with an intra-observer correlation coefficients of 0.99, 0.99, and 0.95 for experienced, trained, and novice observers, respectively (**Figure 5B**). Histologic images from an additional six tissue slides derived from mice with posttraumatic knee OA at 16 weeks post-DMM surgery were scored to validate the reproducibility of the new grading scheme at both the whole joint and tissue-specific levels. Student’s *t*-tests and ANOVA analysis showed no significant differences in averaged OA scores between the observers (**Figure 5C-D**). However, further analysis of the scores for each individual slide revealed discrepancies in OA scores for the subchondral bone change (2/6 slides) and synovitis (2/6 slides) among the Observers, but slide-specific scoring discrepancies for other OA tissues (cartilage, chondro-osteophyte, and peri-articular tissues) were less frequent (0-1/6 slides), suggesting that scoring of the subchondral bone change and synovitis has more variability as it is relatively more subjective than that of other OA tissues, particularly for mild changes. This kind of discrepancy can be resolved by including higher magnification images to help identify more specific histologic features of synovitis, such as proliferation of synovial lining cells and infiltration of immune cells in the subintimal tissue.

**Figure 5.**
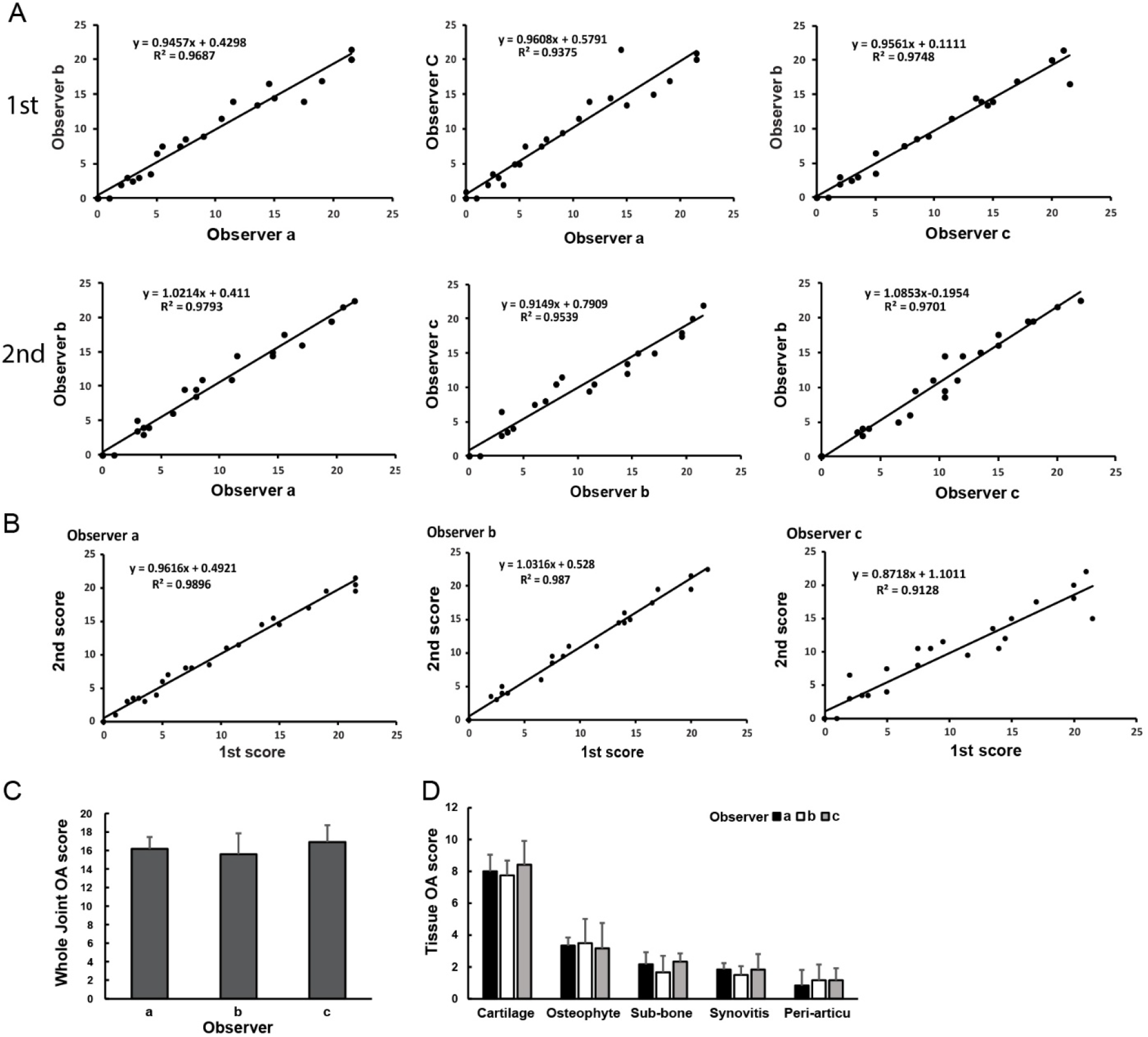
Inter- and intra-observer variation tests showing high reproducibility across experienced and novice observers. (**A**) Correlation coefficients for both first (1st) and second (2nd) measurement scores between the experienced (Observer a), trained (Observer b), and novice (Observer c) (**B**). Intra-observer variability analysis of the first and second measurement scores for each of the three observers. (**C-D**) Averaged OA scores between the observers at the whole joint and tissue-specific levels. Sub-bone = subchondral bone change, Peri-articu = chondrogenesis and endochondral ossification in peri-articular soft tissues.

### Validation of sensitivity and reliability

Histologic images from an additional eighteen tissue slides derived from mice with DMM-induced posttraumatic knee OA at 2, 4, and 16 weeks post-DMM were utilized for sensitivity and reliability assessments.

To test the sensitivity of the new OA grading scheme, averaged whole joint OA scores at 2 and 4 weeks post-DMM were determined by each of the three observers and compared statistically. The data demonstrated that the new grading scheme was able to detect minimal but significant OA progression from 2 weeks to 4 weeks post-DMM (**Figure 6A**). Such sensitivity is critical for determining the efficacy of OA therapies.

**Figure 6.**
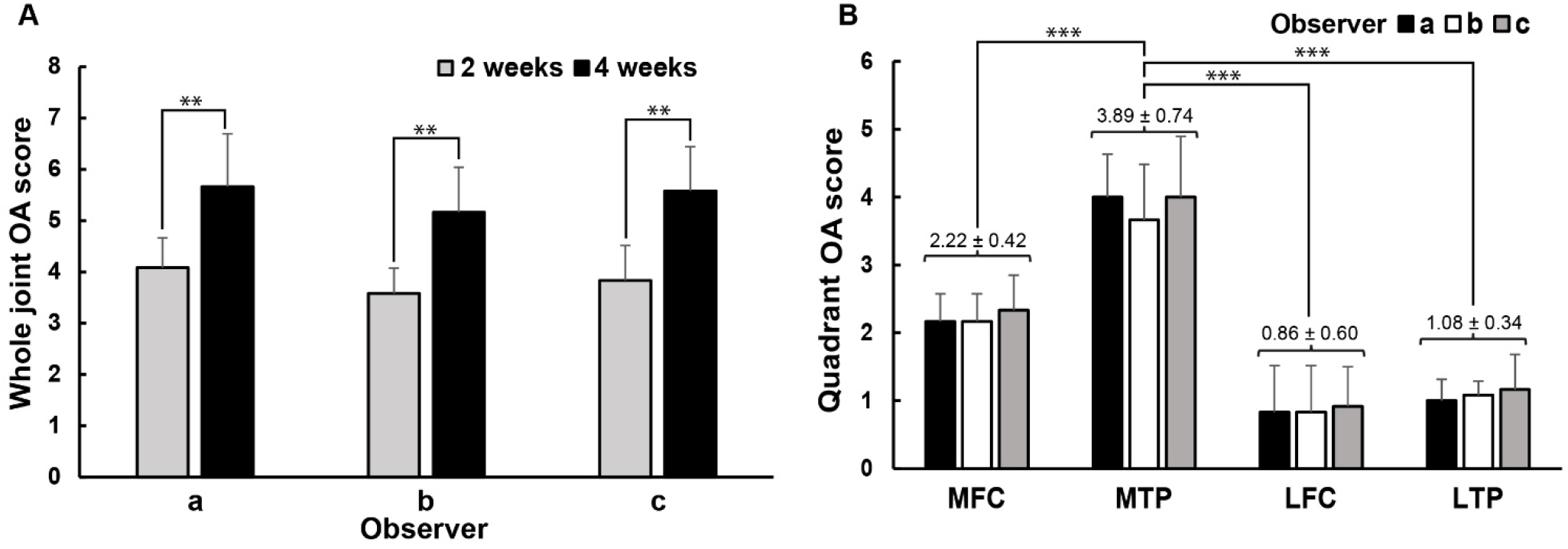
Averaged whole joint scores and quadrant scores showing high sensitivity and reliability of the new OA grading scheme. (**A**) Averaged whole joint OA scores at 2 and 4 weeks post-DMM demonstrate that the new scheme can detect minimal but significant OA progression from 2 weeks to 4 weeks post-DMM. (**B**) Averaged quadrantal OA scores reveal that OA lesion is more severe in the medial tibial plateau (MTP) quadrant than medial femoral condyle (MFC), lateral femoral condyle (LFC) and lateral tibial plateau (LTP) quadrants. The numerical numbers represent mean ± standard deviation from three observers. ** p< 0.01. *** p<0.001 for both A and B.

To test the reliability or accuracy of the new grading scheme, averaged quadrantal OA scores for images of knee OA at 16 weeks post-surgery were determined by each of the three observers and analyzed statistically. As expected, the data demonstrated higher scores (more severe OA) for the medial compartment (MFC and MTP) than the lateral compartment (LFC and LTP), with the most severe OA lesions in the MTP quadrant (**Figure 6B**), consistent with our general histopathologic findings and the reports from others^15; 23^. No differences in OA score were detected between the observers, further confirming the reproducibility of the new grading scheme.

## Discussion

The present study has developed a new comprehensive grading system for murine knee OA based on the authors’ personal experience and extensive literature review. Its inter- and intra-observer reproducibility as well as sensitivity and reliability have been rigorously validated. Although the system was developed using specific mouse models of knee OA, we believe that this new scoring system would be generalizable to other mouse models of knee OA. The advantages, practical applications, and possible limitations of the newly developed scoring system are discussed below.

To score OA severity for the whole joint as an organ, important new OA scoring elements that are frequently seen in osteoarthritic cartilage and non-cartilaginous joint tissues have been added to the new grading system. One of them is the addition of chondrogenesis and endochondral ossification in the peri-articular soft tissues that was not included in any existing OA grading systems. The peri-articular changes are frequently seen in the mouse models of knee OA induced by NFAT1 deficiency, ACLT, and DMM after 16 weeks of surgery, but are not commonly seen in the early phase of DMM-induced knee OA (2-8 weeks after surgery). Additionally, the new system has included a new grade for loss of cartilage surface lamina (grade 1.5), two new grades for severe cartilage loss (Table 1A, cartilage grades 6-7), and a new grade for subchondral bone destruction/cyst formation (Table 1B, subchondral bone grade 2), which were seen in severe spontaneous and posttraumatic OA with a medium- or long-term observation. Therefore, this new grading scheme is suitable not only for mild and moderate OA, but for severe OA seen in late-stage spontaneous and posttraumatic knee OA. Importantly, this proposed grading system includes all major knee joint tissues involved in OA pathogenesis.

To meet the need for scoring large numbers of OA images, simple but reliable and standardized methods for scoring chondro-osteophyte, subchondral bone change, and synovitis are included in the new system (Table 1B). A few previous reports described analysis of multiple joint tissues, but these were non-standardized grading systems that often included additional tests such as micro-CT and/or software-assisted quantitative assessments ^23; 40–42^. The proposed new grading system presented here has the advantage of ease of use without computer software or other equipment items, which is suitable for rapid assessment of multiple joint tissues, particularly when the anatomic location of an OA lesion is unknown.

To increase the sensitivity for detecting early OA changes, grade 1.5 has been added to the new cartilage scoring elements to highlight the loss of surface lamina, which is an important feature of early OA progression. The new scoring system is confirmed to be sensitive to early OA changes and minor disease progression, which is suitable for efficacy evaluations of OA treatments. Standardized and accurate methods of reporting observed OA severity is fundamental to the efficacy assessment of various OA therapies. Without standardization for comparison there is a potential for mis-interpretation of therapeutic results and mis-communication of findings from different study groups.

The proposed grading system looking at essentially all joint tissues known to be involved in the pathophysiology of OA has multiple advantages: 1) understanding the changes in surrounding joint tissues can further elucidate the pathogenic mechanisms of OA; 2) detecting early changes in various joint tissues before cartilage destruction may identify modifiable factors that can be targeted to slow OA progression; 3) understanding if particular models have a higher propensity for changes in synovium versus subchondral bone versus osteophyte formation will further classify individual models and help focus the scope of research.

Weighting of the severity of different tissue types in the grading system is a potential weakness or limitation of this new system as well as the existing OA grading systems. It is recognized that grades 0-2 for chondro-osteophyte formation, subchondral bone change, synovitis, and ectopic endochondral ossification represent a lower “ceiling” score than grades 0-7 for cartilage lesion, which can potentially downplay the involvement of these non-cartilaginous tissues. The proposed scoring system is “semi-quantitative” (no quantitative direct measurements), which may allow for some subjectivity and variability in grading. However, this system allows for relatively rapid grading of slides, increasing the number of slides that can be analyzed per mouse to give a more accurate average. The patellofemoral joint was not included in this analysis, which represents another potential limitation of the grading system, but based on the simplicity of this system it could easily be applied to changes seen within this compartment for specific models. Modifications with respect to the femoral trochlea and patellar facets would have to be established, but the basic principles would remain the same.

## Conclusions

The current study presents a newly developed histologic OA grading system which is modified from but may overcome some weaknesses or limitations of the existing grading systems. The comprehensive grading system presented here covers all-stage osteoarthritic changes in all major OA joint components, including articular cartilage lesion, chondro-osteophyte formation, subchondral bone change, synovitis, and ectopic chondrogenesis and ossification in the peri-articular soft tissues. This modified comprehensive grading system enables us to score OA severity for the whole joint without software-assisted time-consuming measurements, with high intra- and inter-observer reproducibility (correlation coefficients r >0.9) for both experienced and novice scorers. Sensitivity and reliability analyses confirmed the ability of the new system to detect minimal OA progression between two timepoints with a short interval and to identify more severe OA in the medial compartment than the lateral compartment of the knee. This new system may serve as an additional option for OA grading, particularly for rapid assessment of all major joint tissues when the anatomic location of the OA lesion is unknown.

## Acknowledgements

This work was supported by the National Institute of Arthritis and Musculoskeletal and Skin Diseases of the National Institutes of Health (NIH) under Award Number R01 AR059088 (to JW) and the Mary & Paul Harrington Distinguished Professorship Endowment.

## Author Contributions

C. Grote, and J. Wang: Literature search, data acquisition and interpretation, and manuscript preparation. C. Grote, M. Mackay, and J. Wang: OA scoring. X Liu: Statistics and graphic preparation. Q. Lu: Histopathology and selection of representative histologic images. All authors have read and approved the final submitted manuscript.

